# The Transcriptional Program of Regeneration in the Giant Single Cell, *Stentor coeruleus*

**DOI:** 10.1101/240788

**Authors:** Pranidhi Sood, Rebecca McGillivary, Wallace F. Marshall

**Affiliations:** Department of Biochemistry & Biophysics, University of California, San Francisco San Francisco, CA 94158

## Abstract

The giant ciliate *Stentor coeruleus* is a classical model system for studying regeneration and morphogenesis at the level of a single cell. *Stentor* are polarized cells with a complex subcellular architecture. The anterior of the cell is marked by an array of cilia, known as the oral apparatus. This feeding organelle can be induced to shed and regenerate in a series of reproducible morphological steps, previously shown to require transcription. We used RNAseq to assay the dynamic changes in *Stento*’s transcriptome during regeneration with high temporal resolution, allowing us to identify five distinct waves of gene expression. We show that the oral apparatus is a model for organelle regeneration, as well as for centriole assembly and ciliogenesis as many conserved genes involved in those processes are induced. Additionally, we find genes involved in signaling, cell cycle regulation, transcription, and RNA binding to be expressed at distinct stages of organelle regeneration, suggesting that the morphological steps of regeneration are driven by a complex regulatory system.

## Introduction

Regeneration and wound healing are processes that are typically studied at the tissue level in multicellular organisms. A cell scale response to injury is a crucial feature of repair even in multicellular organisms as individual cells also must be able to repair wounds following mechanical disruption. Injured cells must be able to not only patch over the site of injury to prevent leakage of cytoplasm, they also need to re-establish polarity, rebuild organelles and reorganize the cytoskeleton [1]. Recently, deficiencies in cell repair have become implicated in disease, for example in diseases of the heart, lung, and nervous system[2-4]. Yet there is still much to be learned about how an individual cell responds to a wound. A second reason to study regeneration in single cells is to study the mechanisms of how cells build and maintain their shape and organization. Organisms capable of regeneration have long been a focus of developmental biologists, specifically in the area of experimental embryology, as in these systems developmental process can be induced experimentally[5]. Similarly, understanding how cells are able to rebuild cellular components and re-establish global patterning holds the promise of shedding new light on the largely unanswered fundamental question of how cells perform morphogenesis and pattern formation[6,7].

The ciliated Eukaryotic microbe *Stentor coeruleus* is a single cell that can fully regenerate its complex subcellular structure after injury. In this classical system, virtually any portion of the cell, when excised, will give rise to a normally proportioned cell with intact subcellular organization [8,9]. *Stentor* provides a unique opportunity to study regeneration and patterning at the cell scale. Its large size, clear anterior/posterior axis, detailed cortical patterning, and remarkable ability to heal even large wounds in the cell membrane make it especially amenable to surgical manipulation and imaging approaches. Importantly, in *Stentor*, principles of single-cell regeneration can be studied without confounding effects of surrounding cells that may non-autonomously influence an intracellular injury response in the context of tissues. Studies in *Stentor* thus are expected to reveal key general features of wound healing, regeneration, and morphogenesis at the scale of an individual cell.

Transcriptome studies of organisms various multicellular animal species including Zebrafish, axolotl, and Planaria, all of which are capable of life-long regeneration throughout life, have begun to reveal key regulators of regeneration. Many of these studies have delineated the molecular players in regeneration by identifying genes that are expressed when stem cells differentiate into various cell types required to rebuild lost tissue or organs. This transcriptomic approach has thus proven its utility in revealing cell-specific requirements for regeneration in the context of tissues. We have sought to take a similar approach to the problem of single-cell regeneration in *Stentor*.

One of the most dramatic and tractable regeneration paradigms in *Stentor* is the regeneration of the oral apparatus (OA). The oral apparatus is a prominent structure on the anterior side of the cell and is composed of thousands of centrioles and cilia [10] organized into a ciliated ring known as a membranellar band. At one end of the ring is an invagination of the plasma membrane, which is where food particles are ingested. This invagination together with its associated cytoskeletal structures is known as the mouth. The OA can be induced to shed using sucrose shock [11] after which a new OA regenerates over the course of 8 hours, progressing through a series of well-characterized morphological stages (**Figure 1**; [9]). Removal of the macronucleus, at any stage, causes regeneration to halt at the next stage, suggesting that several waves of gene expression may be required to drive different processes at different stages[9]. Chemical inhibitor studies showed that regeneration of the oral apparatus requires transcription [12-15], and it is also known that overall levels of RNA synthesis increase several fold during the regeneration process [13,14,16].

**Figure 1:**
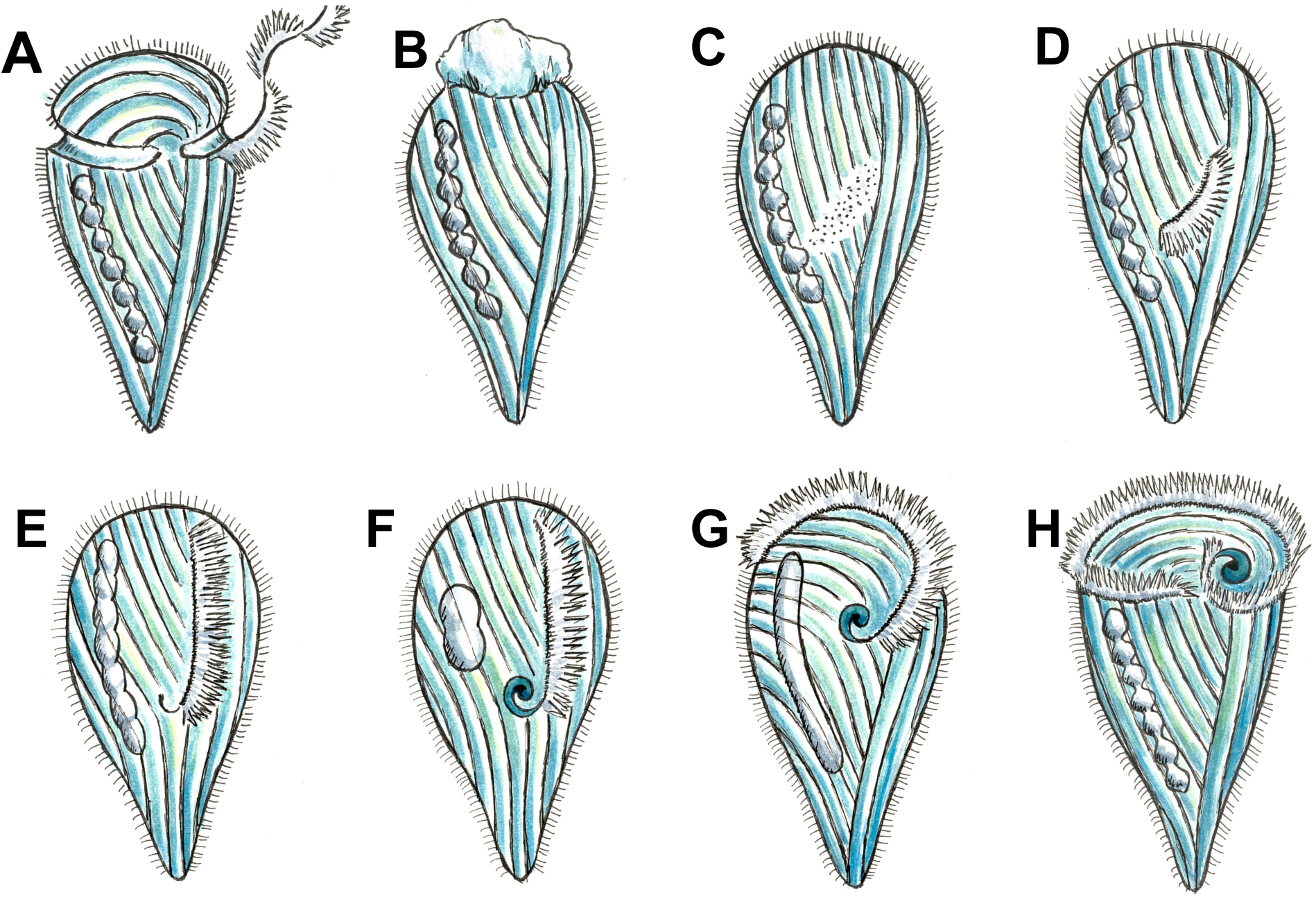
Morphological events in *Stentor* regeneration. (A) The membranellar band is shed during sucrose shock. The body cilia remain on the cell. (B) After the membranellar band is shed, the frontal field protrudes from the anterior end of the cell and is shed. The events in A and B mark the start of regeneration. (C) One hour after the start of regeneration, basal bodies begin to form at the locus of stripe contrast. The blue-green pigment stentorin is cleared from this area. (D) After three hours, the first cilia of the new membranellar band are visible. These cilia show uncoordinated beating. (E) After five hours, the new membranellar band elongates and extends along the anterior-posterior axis. A site for the new mouthparts is cleared at the posterior end of the membranellar band. During this stage, the cilia become oriented with respect to each other and beating becomes synchronous. The nodes of the macronucleus begin to condense. (F) At six hours, the mouthparts are completely formed and the macronucleus is fully condensed. (G) At seven hours the membranellar band and mouth migrate to the anterior end of the cell. The macronucleus extends into a sausage-like shape. (H) By eight hours after sucrose shock the *Stentor* is fully regenerated. The membranellar band completely wraps around the anterior of the cell, the macronucleus is re-nodulated, and the cell resumes normal feeding activity.

Given that transcription is required for regeneration, we hypothesize that there must be a set of genes whose products drive the regenerative process. By learning the identity of these genes, we can determine the molecular pathways involved in building a new OA and coordinating the steps of the process. At the same time, knowledge of the transcriptional program of regeneration would provide a molecular foothold to identify the upstream signals that trigger the process. This type of approach has previously been used successfully to identify genes involved in ciliogenesis, by identify genes are expressed in cells as they regenerate flagella [17,18]. Importantly, these prior studies showed that transcriptomics is a viable approach to studying organelle biogenesis and showed that in addition to identifying structural components of the organelle itself, transcriptomics also reveals genes whose products are not incorporated into the final structure but are required for building it. In this case of a complex regenerative process, this latter strength is of particular importance.

Although *Stentor* regeneration was the focus of study that began over more than 100 years ago and then continuing into the early 1980s, the lack of genetic tools in the organism prevented detailed molecular analysis of its processes. Taking advantage of our recently completed genome of *Stentor coeruleus* [19] and using RNA-seq, we can for the first time provide molecular details of the factors involved in regeneration of the oral apparatus. Key genes which display significant changes in expression over time include highly conserved genes involved in centriole biogenesis and ciliogenesis. Their peak expression corresponds with the timing of morphological stages where these processes occur, confirming that transcriptomic analysis can reveal molecular pathways at work during different parts of the process. Focusing on earlier stages of regeneration, we identified conserved transcriptional regulators as well as RNA binding proteins that are differentially expressed during regeneration. Surprisingly, we find that several highly conserved cell cycle regulators such as Aurora kinases, Rb and E2F, are among the most significantly differentially expressed genes, suggesting a possible role for cell cycle timers in regulating the timing of regeneration. This work opens a new window into the molecular details underlying the century-old question of regeneration in this extraordinary single celled organism.

## Materials and Methods

### Inducing Regeneration and Staging Stentor

Cells were obtained from Carolina Biological Supply and cultured as previously described [20]. Briefly, cells were maintained in Pasteurized Spring Water (Carolina Biological Supply) and fed with Chlamydomonas and wheat seeds. Cells were collected from the same culture for each RNAseq experimental replicate. To induce regeneration, cells were shocked with a 15% sucrose solution for 2 minutes [11], and then washed in Carolina Spring Water thoroughly. Samples of ~20 cells were collected before shock, then at 30 minutes post shock, 1 hour, 2 hours, 3 hours, 4 hours, 5 hours, 6 hours, 7 hours and 8 hours. At each time point, a sample of cells was lysed into RNA-stabilizing buffers specified by the extraction kit, and then stored on ice until the end of the experiment when the RNA purification was performed in parallel on all samples (see below). 4 replicates were analyzed for each time-point.

### Total RNA extraction

RNA was extracted at each time point using the Nucleospin RNA XS kit from Clontech (cat. num. 740902.250). RNA quality was assessed using a NanoDrop and then Bioanalyzer was used to quantify RNA amount. ERCC spike ins (ThermoFisher cat. num. 4456739) were added to each sample in a dilution ranging from 1:1000 to 1:10000 depending on the initial amount of RNA extracted.

### RNA-Seq library preparation and sequencing

RNA-seq libraries were prepared with Ovation RNA-seq system v2 kit (NuGEN). In this method, the total RNA (50 ng or less) is reverse transcribed to synthesize the first-strand cDNA using a combination of random hexamers and a poly-T chimeric primer. The RNA template is then partially degraded by heating and the second strand cDNA is synthesized using DNA polymerase. The double-stranded DNA is then amplified using single primer isothermal amplification (SPIA). SPIA is a linear cDNA amplification process in which RNase H degrades RNA in DNA/RNA heteroduplex at the 5′-end of the double-stranded DNA, after which the SPIA primer binds to the cDNA and the polymerase starts replication at the 3′-end of the primer by displacement of the existing forward strand. Random hexamers are then used to amplify the second-strand cDNA linearly. Finally, libraries from the SPIA amplified cDNA were made using the Ultralow V2 library kit (NuGEN). The RNA-seq libraries were analyzed by Bioanalyzer and quantified by qPCR (KAPA). High-throughput sequencing was done using a HiSeq 2500 instrument (Illumina). Libraries were paired-end sequenced with 100 base reads.

### RNAseq data preparation – trimmed and filtered reads

We used trimmomatic [21] to trim RNAseq reads with the following flags: ILLUMINACLIP:$adapterfile:2:30:10 HEADCROP:6 MINLEN:22 AVGQUAL:20 The settings ensured that we kept reads of at least 22 bases, an average quality score of 20 and trimmed any remaining Illumina adapter sequences.

### Transcriptome generation

To generate a transcriptome, we combined all the reads from all RNAseq samples and timepoints. We ran Tophat2[22] to align the reads to the genome ([19]. We used the following flags to ensure proper mapping in spite of Stentor’s tiny introns: -i 9 -I 101 –min-segment-intron 9 –min-coverage-intron 9 –max-segment-intron 101 –max-coverage-intron 101 -p 20

We then ran Trinity [23] using a genome guided approach. We used the following flags: – genome_guided_max_intron 1000.

### Calculating transcript abundance and differential expression analysis

We used Kallisto to quantify transcript abundance [24] using the following flags: -t 15 -b 30. We then used Sleuth [25] to identify genes which are differentially expressed genes through the regeneration time course. We use an approach similar to that used by Ballgown[26]. Briefly, using a custom script in R, we fit the expression data to time using natural splines (R function “ns”), where the degrees of freedom is 3. Then, using Sleuth, we compared this model to a null model where change in expression is only dependent upon noise. To decide if transcripts were differentially expressed, we defined the minimum significance value (qval in the Sleuth model) to be ten times the minimum significance value of all the ERCC spike-in transcripts. We found that nearly 5583 transcripts are differentially expressed during oral apparatus regeneration. Of these, 485 had no clear homology to proteins in NCBI databases nor PFAM. We identified 234 that did not map to existing gene models. We averaged the expression of all transcripts that mapped to gene models as well as those which were part of a Trinity transcript cluster. All subsequent analysis was performed on these averaged values. Clustering analysis was performed as follows, genes whose maximum expression among the post-shock timepoints was found 30 minutes after sucrose shock were put into one cluster manually. Gene expression profiles before sucrose shock and thirty minutes after are highly correlated (correlation coefficient from Pearson’s correlation = 0.99). The remaining genes were clustered into 4 groups using “clara”.

### Annotation of transcriptome

Following the approach of trinotate (https://trinotate.github.io), we annotated the transcriptome. First we used transdecoder (http://transdecoder.github.io/) to find the longest ORFs (minimum protein length is 100AA and uses the standard genetic code). We used blastx and blastp [27] to search the Uniprot database [28]. Then Hmmscan (hmmer.org, HMMER 3.1b1) was used to search the pfam-a database [29]. Alignments of genes of interest were further manually inspected using a blastp search against the “Model Organism” or “Uniprot-KB/Swiss-Prot” databases.

### Mapping Transcripts to gene models and to genome

We used Gmap [30] to map transcripts to gene models following the approach outlined here: https://github.com/trinityrnaseq/RagonInst_Sept2017_Workshop/wiki/genome_guided_trinity. We used a built in script from Trinity to utilize gmap to align transcripts to a repeat-masked (rmblastn 2.2.27+) Stentor genome. We used bedtools [31] on the resulting bam file to identify overlaps between the aligned transcripts and existing gene models [19].

### Annotation of subsets of genes

We manually curated “ancestral centriole genes” and other genes involved with ciliogenesis and centriole biogenesis [32,33]. We used a reciprocal best Blast search approach to identify genes in the *Stentor* genome with homologs to these manually curated sets of genes.

## Results

### Transcripts are dynamically expressed during regeneration

We performed RNA-seq throughout the regeneration of the oral apparatus (OA) to identify the transcripts involved in *Stentor* regeneration (Figure 2). We focus on regeneration of the OA as it provides an elegant experimental paradigm – a population of cells can be induced to shed their OA synchronously upon simple sucrose shock. The process of building a new OA takes ~8 hours as detailed in Figure 1. RNA samples from ~20 cells were collected prior to sucrose shock, at 30 min after shock, and at 1, 2, 3, 4, 5, 6 7, and 8 hours after sucrose shock. The number of time points was based on the time required to complete regeneration. Taking 10 time points provides a high-resolution time-course of the transcriptional response to regeneration.

**Figure 2.**
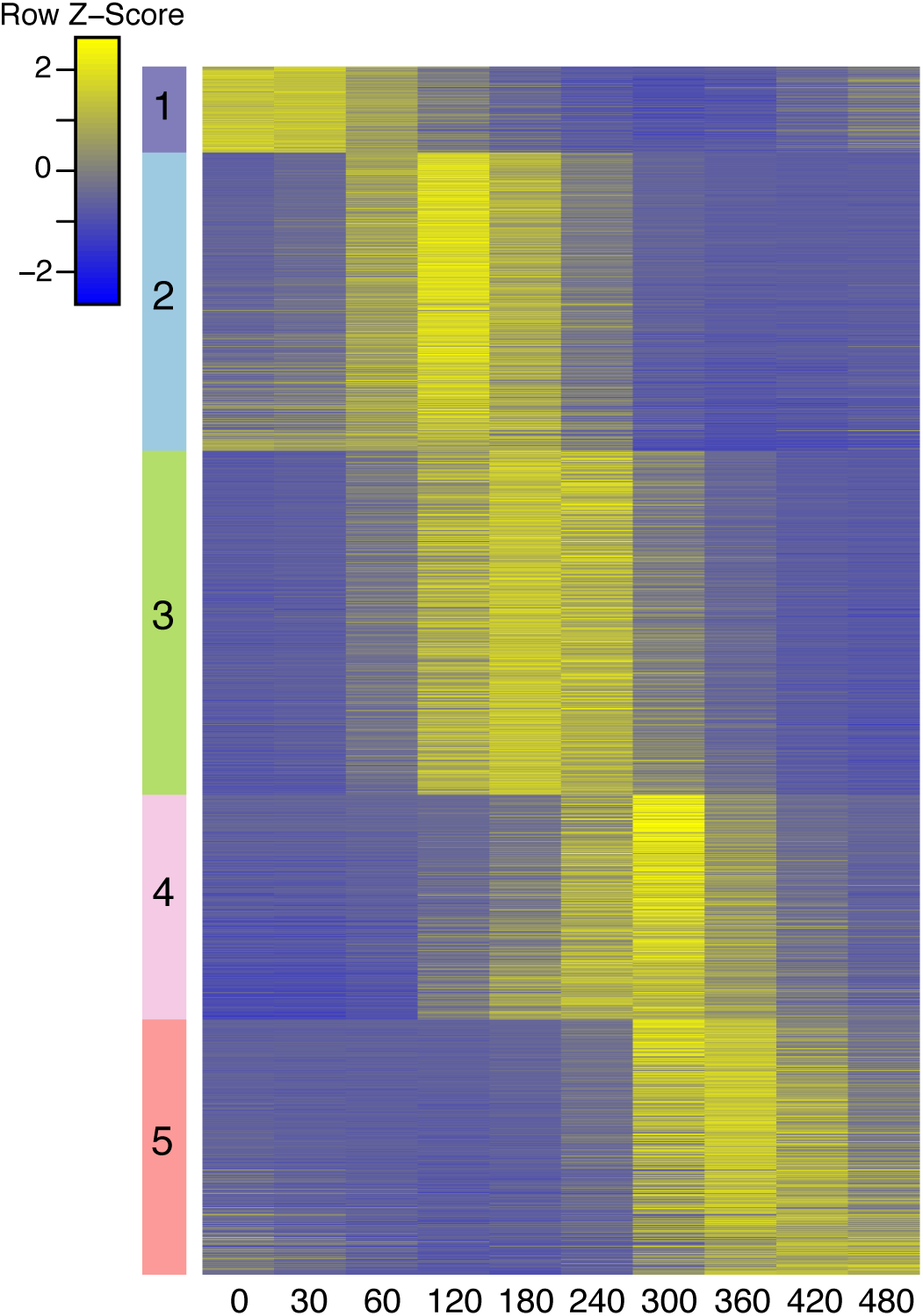
Gene Expression Profiles of Top 3000 Differentially Expressed Genes. Genes are clustered into 5 groups (as indicated by the colored bar on the y axis). Time since sucrose shock (in minutes) is on the x-axis. Group 1 contains all genes whose maximum expression is at the 30 minute time point post sucrose shock. The remaining genes are clustered using “clara”, cluster groups are numbered as indicated and referred to through the text. The peak expression of each cluster of genes corresponds with major development features identified morphologically (Figure 1). Z-score is calculated per row.

At each stage, RNA was extracted, and RNA-seq libraries were sequenced (see Materials and Methods). We combined reads from all samples and replicates and used these to assemble a genome-guided de novo transcriptome[34] using the TRINITY software (RF). We identified 77,638 transcripts in total, of these 65,514 mapped to 30,225 genes. Such a large number of genes is typical of ciliates. Our prior gene prediction for the *Stentor* genome indicated approximately 35,000 genes were present [19].

To identify genes with dynamic expression patterns in response to OA regeneration, we first used Kallisto[24] to map reads to this transcriptome. We used the program Sleuth to identify differentially expressed genes over the time course[25]. Specifically, to identify genes that are differentially expressed we compared two models – a generalized additive model where changes in expression over time are modeled by natural splines, and one in which there is no dependence on time. Sleuth employs a likelihood ratio test that allows for comparison of these models. Overall, we identify 5583 transcripts that exhibited dynamic expression patterns through regeneration but we restricted our analysis to the 3000 most significantly differentially expressed genes, which constitute roughly 10% of the *Stentor* genome. We clustered genes into 5 groups using clara [35] (Figure 2). The first group combines genes whose maximum expression occurs before shock, thus representing genes that are repressed during regeneration, with genes whose expression peaks 30 min after shock. The rest of the genes were clustered into 4 groups whose expression peaked at successively later time periods up to six hours

### Gene Classes

We next focused on specific gene classes to identify modules that may have dedicated roles in OA regeneration. As plotted in Figure 3A, we find that different classes of genes are expressed during various stages of regeneration, suggesting that there are specific molecular requirements for different regeneration steps. The earliest changing genes include transcription factors, consistent with the triggering of a complex transcriptional program. Different transcription factors show altered expression at later stages as well, consistent with the need for ongoing transcriptional regulation. Chaperones are repressed immediately upon induction of regeneration, but then activated in terminal stages. The latest expressed genes include genes encoding components of cGMP signaling (GPCRs and guanylyl cyclases). Kinases are expressed at all stages of regeneration, consistent with the massive expansion of the kinome in Stentor[36] although specific kinase families tend to be expressed at specific stages. One abundant class of kinases observed among the upregulated genes was the dual-specificity DYRK kinases, with 17 different DYRK family members expressed among several different expression groups. However, the kinome of *Stentor coeruleus* has been predicted to contain 142 DYRK family members, making them one of the most highly expanded kinase families in the genome [36]. Given that the *Stentor* genome contains 35,000 genes, we would expect that roughly 14 DYRK would be present in any randomly chosen set of 3,000 genes. As such, DYRK genes are not particularly enriched among the differentially expressed genes during regeneration. Genes involved in centriole assembly are, on average, expressed earlier than genes involved in subsequent formation of cilia by those centrioles. Genes involved in conversion of the centrioles to basal bodies are expressed prior to formation of cilia. Among the classes of cilia-related genes examined, those involved in ciliary motility are, on average, expressed later than those involved in ciliary assembly. These observations are consistent with a sequential gene expression program driving the stepwise assembly of motile cilia. Finally, OA regeneration involves major dynamic changes to the cytoskeleton of the cell. Kinesin motor proteins, which drive these types of changes, are expressed consistently throughout the process, with a major induction of these proteins early in the process (in the third clustered group). Four specific gene classes are plotted in Figure 3B-E, which further illustrates that different genes and gene families display specific kinetics of expression throughout the process of regeneration. We describe below additional membership of each of the cluster of genes. Notably, many genes from all 5 clusters also have orthologs conserved in mammals.

**Figure 3.**
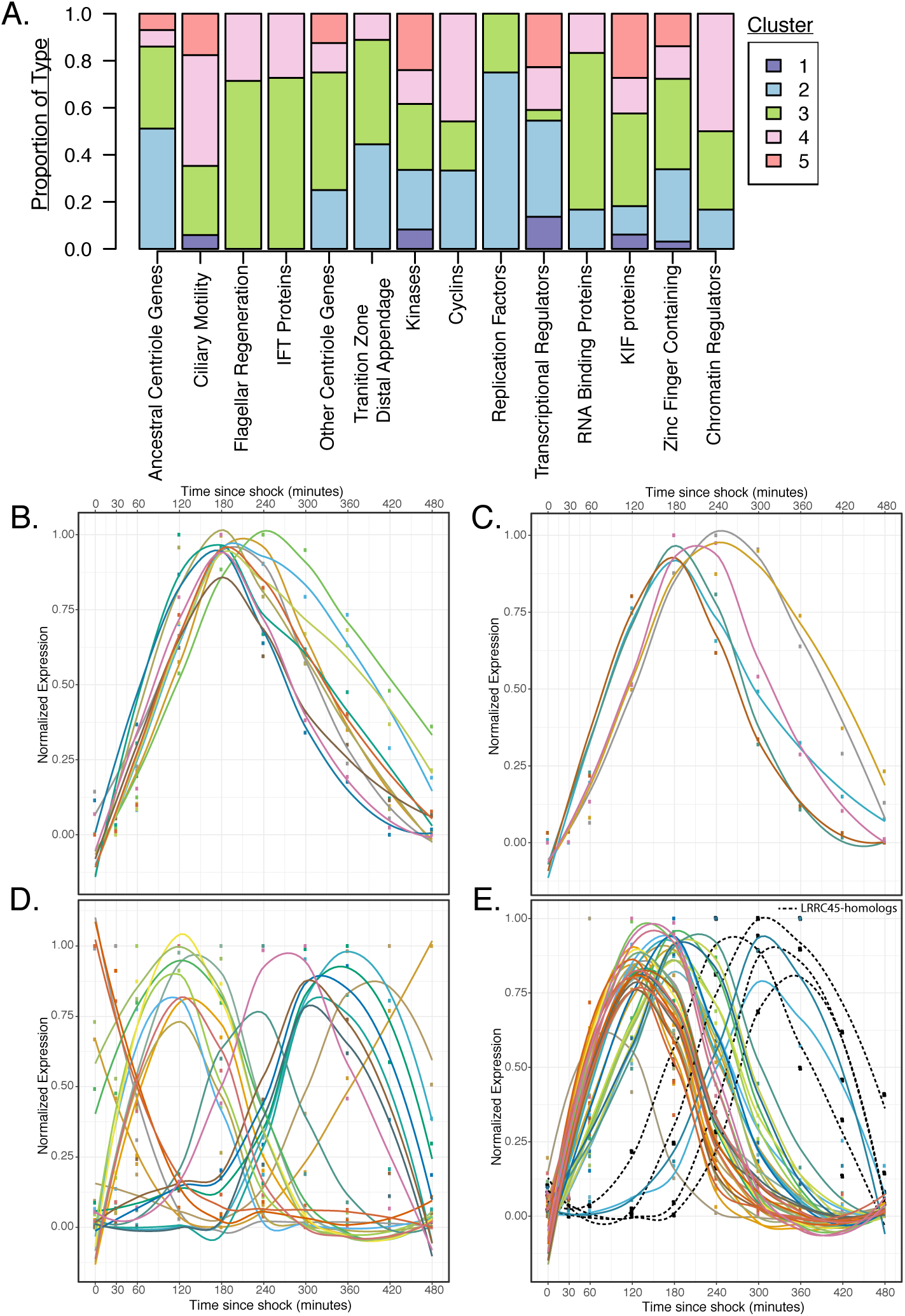
Functional Annotations of top 3000 differentially expressed genes. A) Stacked barplot of the cluster membership for genes of various categories. Number of genes in each category (Zinc Finger Containing – 65; KIF proteins – 33; RNA Binding Proteins – 6; Chaperones – 3; Transcriptional Regulators – 22;Replication Factors – 8; Adenylyl and Guanylyl Cyclases – 3;GPCR – 2;Cyclins – 24; Kinases – 146; Cilia-related Transition Zone Distal Appendages – 9; Other Centriole Genes – 8; IFT Proteins (cilia-related) – 11; Flagellar Regeneration (cilia related) – 7; ciliary motility – 17; ancestral centriole genes – 43). B) Expression profiles for genes involved in intraflagellar transport (IFT) C) Expression profiles for genes orthologous to genes upregulated during flagellar regeneration in Chlamydomonas D) Expression profiles for transcriptional regulators E) Expression profiles for “Ancestral Centriole Genes” – LRRC45 homologs are indicated by black dashed lines. Normalized Expression plotted in panels B-E is calculated by linearly mapping expression levels between the minimum and maximum level measured for each gene during the regeneration timecourse. Curves are loess fits over points.

#### Group 1

The patterns of gene expression that we observe directly correlate with the known morphological changes taking place during OA regeneration (Figure 2). Group 1, contains 213 genes which are immediately repressed upon induction of OA regeneration. Among these are three transcriptional regulators – SFL1-like, HSF, and a myb-like transcription factor. In other systems, both SFL1-like and HSF regulate stress responses[37]. Another quickly repressed protein has homology to the splicing factor YJU2 which is an essential protein required for mRNA splicing in yeast[38]. Other proteins found in this first group are Zinc-finger containing proteins, which may be involved in transcriptional regulation through DNA binding activity, kinesins, and a DNAJ domain containing protein. There are many kinases found in this group, but key among these is a homolog of Aurora kinase (see Figure 4D for more details about Aurora kinase dynamics during regeneration). Finally, a homolog to RSP1 is also among the group of immediately repressed genes[39].

**Figure 4.**
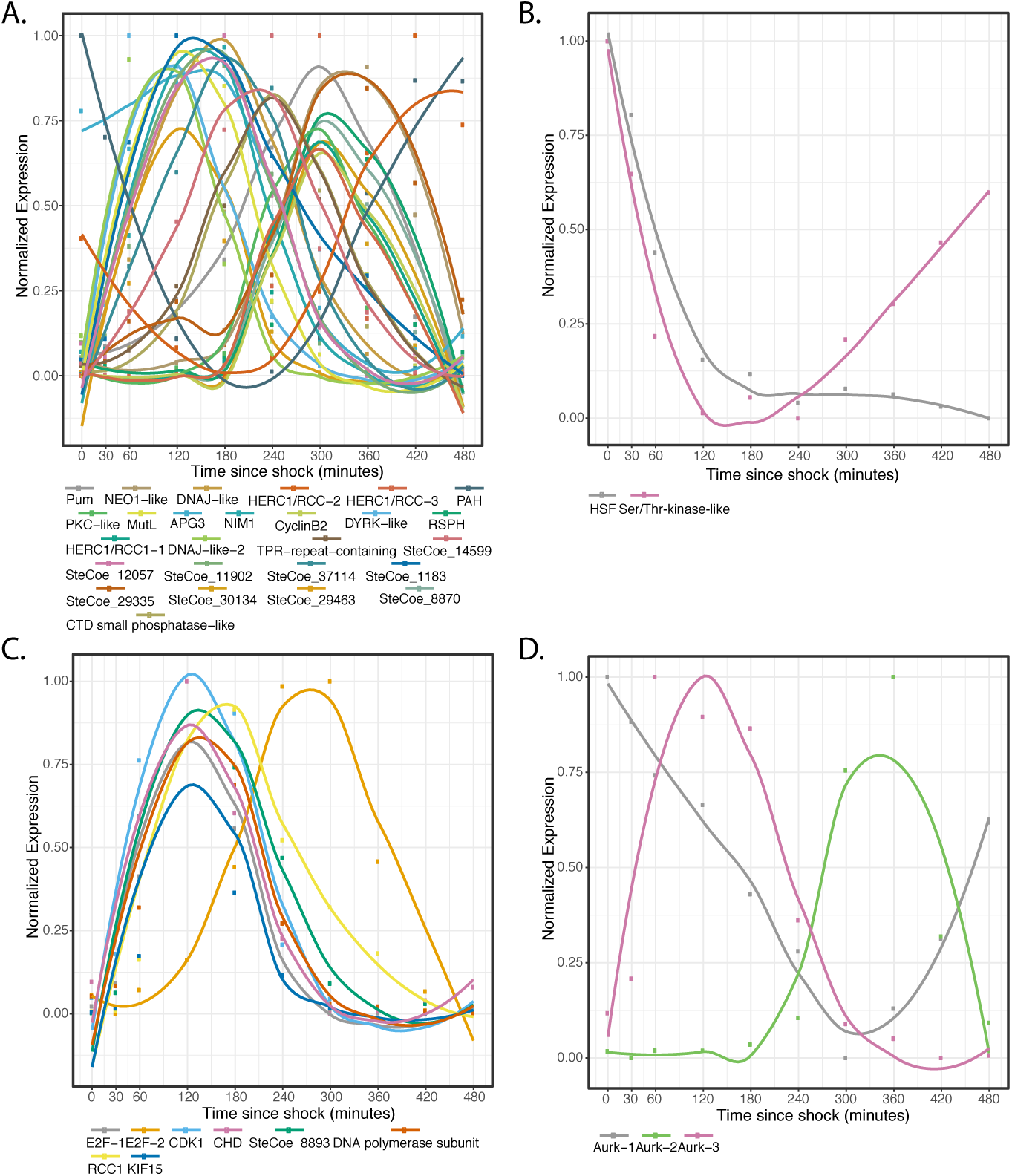
Gene Expression Profiles of putative regulatory genes and their targets. Curves are loess fits. Normalized Expression plotted in all panels is calculated by linearly mapping expression levels between the minimum and maximum level measured for each gene during the regeneration time course. (A) Predicted Targets of the RNA-binding protein, Pumilio (B) Predicted Targets of HSF (C) Predicted Targets of Transcriptional Regulator E2F (D) Aurora Kinase homologs.

#### Group 2

Group 2 contains 741 genes, most of which reach peak expression an hour after sucrose shock. Among these is a homolog of YPT1, a Rab-family GTPase that in other systems is involved in mediating the progression of Golgi cisternae [40]. We also find a SET1-domain-containing protein with histone methyltransferase activity, as well as an RRM-1 domain containing protein with predicted RNA binding activity. Additionally, this group is enriched for replication factors. Among this group we find homologs to replication licensing factors MCMs [41] and RFC-like proteins [42], in addition to a component of the DNA Polymerase a primase which is required for initiation of replication[43]. Another exciting finding is that many known regulators of the cell cycle are induced in this group of genes. First, multiple homologs of the Rb transcriptional regulator are expressed in this group –4 of the 6 homologs found to be differentially expressed during regeneration. In this group, we also find one of the two E2F homologs differentially expressed during regeneration. Together, Rb and E2F regulate the transition from G1-S in other systems[44]. Additionally, another Aurora kinase is found to be among this group. In other systems Rb, E2F and Aurora kinases interact to prevent endoreduplication during the cell cycle [45]. The large number of conserved replication factors and cell cycle genes found in this early induced group suggests a link between these processes and regeneration.

The time of maximum expression of group 2 genes corresponds to the time in regeneration at which thousands of centrioles assemble de novo to form a so-called “anarchic field” [46]. These centrioles will ultimately organize themselves into arrays and become the basal bodies that nucleate the ciliature of the OA. If RNAseq truly reflects the molecular events of regeneration, we would expect to see strong upregulation of genes known to be involved in centriole biogenesis. We therefore analyzed our data for the presence of a set of genes known as the “ancestral centriolar genes” found in any organism that has a canonical centriole [47]. Among these are sas6, sas4, poc1,cetn3, cetn2, tbc31, rttn, f161, cep76, cep135, cep120, ccdc77, and ccdc61[33]. SAS6 is notable as one of the earliest known factors involved in assembling the ninefold symmetric structure of the centriole. Out of the 29 most conserved ancestral centriole genes, we find that 22 are expressed in either group 2 or group 3. This expression of the core centriole gene set at the exact stage when centrioles are forming thus provides a biological confirmation of our analysis.

#### Group 3

Group 3 contains 854 genes. As with group 2, group 3 contains genes involved in centriole biogenesis (sas6, poc1, ofd1, lrrc9, cetn3, cetn2, cep41, cep164, cep135, cep131, tri37). Notably, Cep164 is involved in assembling the distal end of the basal body where it attaches to the cortex, and TRI37 is involved in terminating centriole duplication [48]. Group 3 also contains many key proteins involved in ciliary biogenesis. Among these are Intraflagellar transport (IFT) proteins, which are required for transport of protein in assembling cilia, components of the BBsome–another complex involved in ciliary assembly–as well as genes whose expression is known to increase during flagellar regeneration in Chlamydomonas [17] and which are involved in ciliogenesis in other species, such as MNS1 and MIPT3. This time point corresponds to the time in regeneration at which the cilia begin to assemble on the newly formed basal bodies [10]. This group also contains the most proteins from the kinesin super family (13 of the 24 found to be differentially expressed).

Additionally, we find that a homolog to the splicing factor CEF1 is expressed at this time point, as are two putative chromatin regulators with histone acetyltransferase domains. We also find putative homologs to the RNA binding proteins, piwi and rae-1. In other ciliates, Piwi proteins have been found to play two roles: one is for mediating RNAi during vegetative growth and the other is for mediating reorganization of the micronuclear genome during mating[49,50]. In other systems, RAE1 binds mRNAs and transports them from the nucleus to the cytoplasm by associating with microtubules[51].

#### Group 4

Of the 558 genes in group 4, we find chromatin regulators including a histone deacetylase and two histone lysine methyltransferases. At this timepoint, the macronucleus begins to undergo drastic morphological changes, similar to those seen during cell division. This is also the group that contains the most cyclins. Additionally, this group contains a major wave of expression of genes involved in regulating ciliary motility including spag6, several radial spoke genes, and components of the dynein regulatory complex. None of these genes is required for the assembly of cilia in other model systems, but instead are involved in coordinating the activity of axonemal dyneins to generate motility [62]. In *Chlamydomonas*, radial spoke protein synthesis reaches its maximum rate 30-60 minutes after the flagella have begun assembling [52], which roughly matches the delay seen in our data between genes involved in ciliary assembly and genes encoding radial spokes during *Stentor* regeneration. The timing of group 4 also correlates with the time period during which the oral cilia, which initially undergo random beating, begin their characteristic coordinated beating motility to form metachronal waves [10].

#### Group 5

Group 5 contains 634 genes which are the last to reach peak expression during regeneration. Among these are kinases, kinesins and zinc finger containing proteins. We also find myb-like transcriptional regulators and Rb-homologs. Interestingly, the only clear centriole-related genes we find in this late-expressed group are three homologs to lrrc45 which is a linker component required for centriole cohesion[53] (a fourth is found in group 4). In this regard, we note that when centrioles first assemble during oral regeneration, they do so with random orientations relative to each other, creating a so-called “anarchic field” [46]. It is only later in the process that the centrioles associate into pairs and then larger groups to form the membranelles that are the dominant ultrastructural motif of the oral apparatus. The expression of lrrc45 at exactly this stage suggests that this linker may be a key element for assembling the membranellar band from the initially randomly oriented centrioles.

### Data validation: comparison with expected features of expression program

Several features emerge from the analysis of gene clustering during regeneration which match our *a priori* expectations about the gene expression program of OA regeneration and thereby serve to confirm the validity of our results. First, the duration of each gene expression group is roughly one to two hours, corresponding to the length of time that regeneration is known to proceed following surgical removal of the nucleus [9]. The persistence of regeneration over this time scale likely reflects the lifetime of the mRNAs that drive each stage. This time scale also matched the period of time during which visibly distinct morphological processes occur, for example ciliogenesis initiates in different regions of the oral primordium at slightly different times, with early-stage events of ciliogenesis taking place over a roughly 1-2 hour period [10]. Therefore, we expected that groups of related genes would show peaks of expression lasting on the order of 1-2 hours, as we observed. Second, the number and timing of the five clusters correspond with the number and timing of known morphological events, consistent with our *a priori* expectation that different morphological events in regeneration may be coordinated by distinct modules of genes. Finally, we observed a strong correlation between the types of genes expressed at a given stage, and the cell biological events taking place at that stage. Based on this correlation, we believe that examination of other genes with correlated expression patterns will reveal previously unknown molecular players in organelle regeneration, centriole biogenesis and ciliogenesis.

### Transcriptional Regulation during Regeneration

What drives the timing of the steps of regeneration? Given that regeneration entails a complex program of gene expression, we hypothesized that different transcription factors might trigger genes at different stages of the process. Our clustering analysis revealed an E2F transcription factor expressed at the two-hour time point. Based on this result, we identified putative E2F targets based on promotor motif analysis [54], and asked whether these predicted targets exhibited specific expression patterns during regeneration. As shown Figure 4C, there is indeed a tight pattern of E2F targets expressed during a one-hour window that corresponds roughly to the time in regeneration at which centriole related genes are expressed (Figure 3C). This pattern closely matches the pattern of expression of the E2F ortholog, designated E2F-1. A second E2F ortholog is expressed much later in regeneration as designated by the orange curve in Figure 4C. This analysis suggests that E2F-1 may be a driver of early expression patterns. In contrast to E2F which is upregulated during regeneration, the HSF transcription factor, involved in stress response, is downregulated during regeneration (Figure 4B). Motif analysis of potential HSF targets [55] revealed a Serine/Threonine kinase whose expression also decreased during regeneration, with similar kinetics as HSF itself, but then restored activity at later stages of regeneration.

In addition to transcription factors, we also noticed that the RNA binding protein Pumilio was upregulated during regeneration. Given the importance of Pumilio for mRNA localization and translation control during pattern formation in embryos such as *Drosophila*[56-58], which is of a similar size scale as *Stentor*, we hypothesized that this factor may play a role in regulating key regeneration regulatory factors at a post-transcriptional level. If this were true, then we would expect the gene expression program of regeneration to include genes whose messages contain Pumilio binding sites. Analysis of Pumilio recognition motifs [59] in the 3’ UTRs of differentially expressed genes confirmed this expectation (Figure 4A). There is a tendency of the putative Pumilio targets to have two peaks of expression at 120 minutes and 300 minutes after the start of regeneration, though there are predicted targets expressed throughout the stages of regeneration. Of the 25 predicted targets, the 16 with recognizable homology correspond to a wide variety of protein types, including DRYK and PKC family kinases, as well as phosphatases, cyclins, and DNAJ domain proteins, suggesting a global effect of Pumilio on many aspects of the process of re-patterning the cell.

### Cell cycle regulators during regeneration

A final class of regulatory genes we consider here are cell cycle regulatory kinases. The morphological steps of OA regeneration visible on the cell surface (Figure 1) are virtually identical to the steps by which a new OA forms during normal cell division [9]. A hint at a connection between regeneration and cell division is also provided by morphological changes that take place in the nucleus. Like other ciliates, *Stentor* contains a single large polyploid macronucleus that contains up to a million copies of the expressed genome [19], as well as several smaller diploid micronuclei. During division, the micronuclei undergo spindle-based mitosis, but the macronucleus does not. Instead, it is simply pinched in half by the cleavage furrow. Prior to this pinching, the elongated macronucleus shortens and compacts into a more spheroidal shape, which then re-elongates just before cytokinesis. Interestingly, these same shape changes occur during regeneration, even though the cell is not going to divide [9,60]. The strong morphological similarities between regeneration and division, at both the cortical and nuclear level, suggest that OA regeneration might involve co-option of parts of the cell cycle machinery to regulate the timing of events. Indeed, the ability of the cell cycle machinery to regulate sequential events is one of its hallmark features. Consistent with this prediction, we have found that numerous cell cycle related genes are upregulated during regeneration. These include homologs of E2F, Rb, and various cyclins. Previous analysis of the *Stentor* kinome identified 44 Aurora related kinases [36], of which three are found to be differentially expressed during regeneration. Figure 4D plots the expression pattern of these three Aurora kinases as a function of time during regeneration. Two (Aurk2 and Aurk3) are induced during regeneration, but at distinct time points, with Aurk3 peaking at one hour after the start of regeneration, when centrioles and cilia are forming, and Aurk2 peaking at four hours, when the membranellar band of the OA is beginning its anterior migration. A third Aurora kinase, Aurk1, also shows differential expression during regeneration but in this case, it is repressed once regeneration begins and reaches a minimum around five hours after the start of the process, before eventually returning to its pre-shock level. The three curves in Figure 4D suggest an inflection point at approximately four hours, when Aurk2 levels are rising, but Aurk1 and Aurk3 levels are decreasing.

## Discussion

Our analysis of transcription during oral apparatus regeneration in *Stentor coeruleus* reveals key pathways and genes that are involved in the regeneration and re-patterning of a single cell. We find tight temporal correlation between the expression of centriole and cilia related genes and the corresponding events of centriole biogenesis and ciliary assembly, respectively. Based on this positive result, we predict that at least some of the genes in these clusters with no or poor homology to known genes may encode undiscovered factors involved in centriole biogenesis and ciliogenesis. While the proteome of the centriole is by now well characterized, we hypothesize that cluster 2 may contain genes whose products are needed for centriole assembly, but may not encode structural components of the centriole itself.

One particularly notable result from our analysis is the expression of many cell division and cell cycle related genes, particularly in the later stages of regeneration. We hypothesize that these expression patterns may reflect a mechanistic role for the cell cycle machinery in regulating the timing of regeneration. This potential connection highlights a classical question in the biology of regeneration: is regeneration a distinct process in its own right, or instead does it reflect a reactivation of development? In the case of *Stentor*, our results support the latter view.

The regeneration of the *Stentor* oral apparatus is but one of many regeneration paradigms that have been described in this remarkable organism. Another striking example of regeneration is the ability of *Stentor* to regenerate two intact cells after a single cell is surgically cut in half. The posterior half cell has to regenerate an OA, while the anterior half cell has to regenerate a tail. A transcriptomic analysis of bisected *Stentor* regeneration has just been reported in the related species *Stentor polymorphus[61]*. It will be interesting to ask to what extent the posterior cell expression pattern in their study reflects the OA expression pattern in our study.

More than a century ago, microsurgical studies in *Stentor coeruleus* were performed by leading developmental biologists, including Morgan, Lillie, and Balbiani. Although these studies revealed great detail about morphological changes during regeneration, the organism was never developed as a molecular model system and thus, apart from inhibitor studies, there has been little information about the molecular basis of regeneration in *Stentor*. Genomic approaches in *Stentor* are now feasible and, as described here, we have begun to reveal key molecular details of intracellular patterning and regeneration mechanisms. Already, we find that these many of these programs are conserved across Eukaryotes. As such, further delineation of these programs holds great potential for the discovery of therapeutics to aid in cell recovery from injury as well as for providing a way to identify conserved molecules involved in the origins of cell geometry.

## Acknowledgments

This work was supported by an American Cancer Society postdoctoral fellowship (PS) and NIH grant R01 GM113602 (WFM). Library preparation and QCs for sequencing was conducted by the Gladstone Institute Genomics Core. The authors thank members of the Marshall lab for many helpful discussions.

